# Motifs of human hippocampal and cortical high frequency oscillations structure processing and memory of naturalistic stimuli

**DOI:** 10.1101/2024.10.08.617305

**Authors:** Akash Mishra, Gelana Tostaeva, Maximilian Nentwich, Elizabeth Espinal, Noah Markowitz, Jalen Winfield, Elizabeth Freund, Sabina Gherman, Ashesh D. Mehta, Stephan Bickel

## Abstract

The discrete events of our narrative experience are organized by the neural substrate that underlies episodic memory. This narrative process is segmented into discrete units by event boundaries. This permits a replay process that acts to consolidate each event into a narrative memory. High frequency oscillations (HFOs) are a potential mechanism for synchronizing neural activity during these processes. Here, we use intracranial recordings from participants viewing and freely recalling a naturalistic stimulus. We show that hippocampal HFOs increase following event boundaries and that coincident hippocampal-cortical HFOs (co-HFOs) occur in cortical regions previously shown to underlie event segmentation (inferior parietal, precuneus, lateral occipital, inferior frontal cortices). We also show that event-specific patterns of co-HFOs that occur during event viewing re-occur following the subsequent three event boundaries (in decaying fashion) and also during recall. This is consistent with models that support replay as a mechanism for memory consolidation. Hence, HFOs may coordinate activity across brain regions serving widespread event segmentation, encode naturalistic memory, and bind representations to assemble memory of a coherent, continuous experience.

## Introduction

Despite our daily immersion in continuous, multimodal experiences, we tend to recall memories as episodic events^1^. These “real life” narratives facilitate connections with other experienced events to generate memory scaffolds^2^. We have an insufficient understanding of how the human brain segments a continuous narrative, coordinates the activation of widespread memory-containing cortical regions, and integrates events experienced over time^3–5^. It is likely that these processes rely on a synchronizing mechanism that organizes several brain regions, including the hippocampus, sensory cortices, and widespread cortical regions that process and store multimodal memories.

A continuous, naturalistic narrative may be divided into discrete events by event boundaries, or timepoints that are associated with contextual shifts^6,7^. This process is supported by an event segmentation hierarchy that spans lower-level sensory regions and higher-order, multimodal processing regions^8,9^. Critically, different levels of this assembly are receptive to varying degrees of event granularity: sensory regions are more responsive to finer boundaries whereas multimodal integration regions are more responsive to abstract, coarse boundaries^8,10,11^. The critical process of recapitulating event representations from previously-experienced events, or “replay,” occurs following event boundaries; hence, this may represent an optimal window for the consolidation of continuous stimuli memory^7,12^. Several studies have suggested that the hippocampus may play a role in coordinating event segmentation and memory processes during this window^7,8,12–16^. However, the mechanisms underlying this are not well understood.

80-140Hz high frequency oscillations (HFOs) have been implicated in the recruitment of memory-containing cortical regions^17–22^, especially during the replay of previously-experienced events^23,24^. HFOs that occur at the same time in different brain regions (“co-HFOs”) may synchronize brain regions in the memory network^19,21,25–28^ by eliciting specific patterns of neuronal firing^20^. Since the process of naturalistic event memory likely requires widespread, brain-wide coordination, HFOs may serve as an optimal potential coordinating mechanism underlying this process. We hypothesize that co-HFOs bind hippocampal-cortical activity in cortical regions previously found in fMRI studies to be involved in event segmentation (for example, angular gyrus, precuneus, lateral occipital cortex)^8,29^. Further, based on scalp EEG^30,31^ and fMRI^29^ studies showing pattern similarity between event viewing and following event boundaries, we expect co-HFOs to exhibit specific spatial and temporal patterns (“co-HFO motifs”) that reveal similarity in the processes of event encoding (viewing), replay (following event boundaries), and memory retrieval (recall). As ensembles of micro- and meso-scale HFOs may serve as engrams for memory processes^32^, these co-HFO motifs would be unique for each event. In addition, since memory processes scaffold upon several previously-viewed events, they would contain representations for both the most recently-viewed event and several events prior.

In this study, we recorded intracranial electrophysiological data from patients who watched a 10- minute clip of the cartoon *Despicable Me* and then performed a free recall of their event memory. We leverage the spatial and temporal resolution of intracranial electroencephalography to show that hippocampal HFOs increase in rate following event boundaries, and this activation is most prominent in right anterior hippocampal electrodes. Hippocampal-cortical co-HFOs increase in a wide cortical network, with the most prominent increases present in regions that show event boundary activation in prior fMRI studies^8,29^. We identify event-specific spatial and temporal patterns of HFOs (“co-HFO motifs”) that recur following event boundaries and during event free recall. The magnitude of pattern reactivation for an event diminishes as additional events are viewed. Taken together, HFOs represent a potential mechanism of synchronizing activity across the expansive hippocampal-cortical memory network and may underlie how humans process continuous, real-life stimuli as a sequence of events. This process may be associated with event-specific arrangements of co-HFOs that are present during event viewing and reactivate at times of memory function.

## Results

The aim of this study was to investigate HFOs as a mechanism underlying multi-regional synchrony at event boundaries. To answer this, we recorded intracranial electroencephalography (iEEG) from 32 testing sessions (30 patients) while they watched a 10-minute clip of the film *Despicable Me* (Figure 1A). Nine event boundaries with high inter-subject consistency^9^ were used. Twelve subjects were additionally asked to freely recall details from the clip. These subjects recalled an average of 55.8±17.8% of events (Figure 1B).

**Figure 1.**
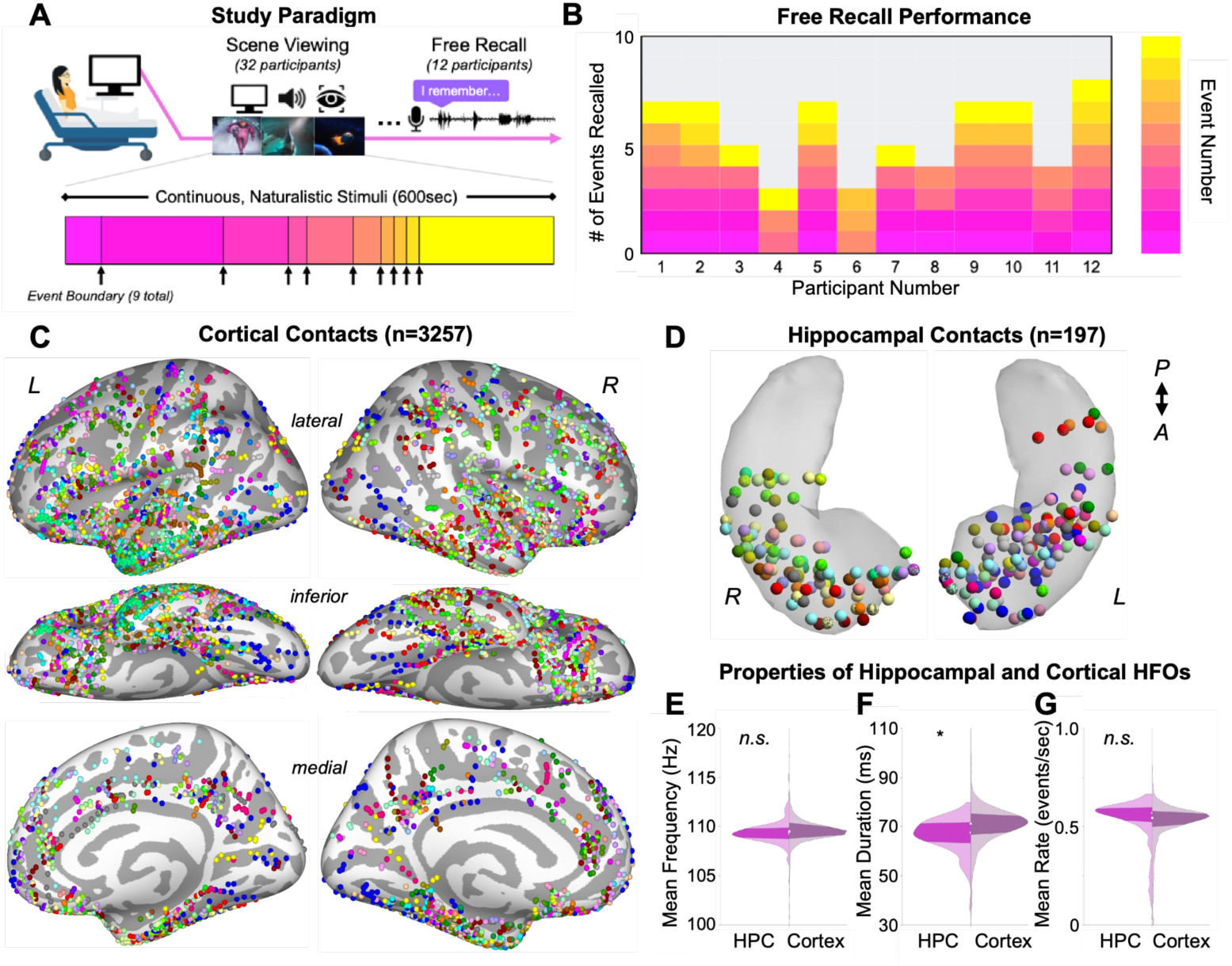
Experimental design, recording contacts, and HFO detection. (A) Experimental design. Subjects (n=32) viewed a 10-minute continuous clip of the film *Despicable Me*. The clip was segmented into discrete 10 events by 9 event boundaries. Following this, a subset of subjects (n=12) performed a free recall describing what they remembered from the clip. (B) Chart detailing content of recall for all 12 subjects who performed free recall. (C) Cortical contact locations across subjects on a standard inflated brain. Each color represents one subject. (D) Locations of hippocampal contacts across subjects on a hippocampal surface of a standard brain. Each color represents contacts from individual subjects. Distribution of hippocampal and cortical HFO (E) peak frequency; (F) duration; and (G) rate across n=3257 cortical contacts and n=197 hippocampal contacts. * denotes significant difference of means at p<0.05 level (student’s t-test); n.s., not significant.

### Properties of hippocampal and cortical HFOs

We examined HFOs during scene viewing (3257 cortical and 197 hippocampal contacts across 32 sessions) (Figure 1C-1D) and free recall (1075 cortical and 90 hippocampal contacts across 12 sessions) (Figure S1). Detected HFO events during viewing in hippocampal and cortical contacts did not differ significantly in peak frequency (mean 109.42±0.07Hz in hippocampus and 109.52±0.02Hz in cortex; t(3465)=1.119, p=0.263; Figure 1F) or rate (mean 0.531±0.008 events/sec in hippocampus and 0.522±0.002 events/sec in cortex; t(3465)=1.234, p=0.216; Figure 1H), but cortical HFOs were of a longer duration (mean 66.68±0.43ms in hippocampus and 70.22±0.14ms in cortex; t(3465)=6.711, p<0.001; Figure 1G). Cortical HFO characteristics are stable across cortical regions (Figure S2).

Dotted line represents the mean hippocampal HFO rate over this epoch. (B) Magnitude of change in hippocampal HFO rate following event boundaries (compared to before the boundary) for each hippocampal contact on a normalized hippocampal surface. Warmer colors indicate greater increase in hippocampal HFO rate. (C) t-values obtained from a generalized linear model assessing characteristics of hippocampal contacts that are associated with increases in hippocampal HFO rate following event boundaries. Factors tested were: longitudinal axis (‘ant’), hemisphere (‘hem’), and CA1 subfield (‘CA1’). * denotes statistical significance at p<0.05 for the model term or interaction effect. (D) Mean change in hippocampal HFO rate following event boundaries (compared to before boundaries) for hippocampal contacts, divided by hemisphere and longitudinal axis of the hippocampus. Error bars represent one standard error of the mean. * denotes statistical significance at p<0.05 (Bonferroni-corrected; one-sample t-test).

### Hippocampal HFOs during scene viewing and recall

Event boundaries may serve as a period of increased hippocampal-cortical coordination for the consolidation of previously-viewed events. Hence, we examined the relationship between hippocampal HFOs and event boundaries. We constructed an event boundary-locked peri-event time histogram (PETH) across all subjects and hippocampal contacts. This was compared to a PETH derived from visually- and auditory-matched control scenes (Figure S3). Hippocampal HFO rate increased transiently 1 to 2 seconds following event boundaries (p=0.026, cluster-based permutation test; Figure 2A; subject-level analysis: paired t(31)=1.779, p=0.085; Figure S4). For subjects who performed free recall, scenes that were subsequently recalled demonstrated increased hippocampal HFO rate in this post-event boundary window compared to scenes that were not subsequently recalled (p<0.07; Figure S5). This supports the hypothesis that hippocampal HFOs play a role in replay processes in the period following event boundaries^33^. Next, we aimed to assess which hippocampal contacts contribute to this phenomenon. We calculated the magnitude of HFO rate increase following event boundaries for each contact (2- second window following event boundaries compared to 2-second window prior) (Figure 2B). Then, we implemented a general linear regression model with the following factors: hippocampal subfield (CA1 or non-CA1), longitudinal position along the hippocampus (median split by patient; anterior or posterior), and brain hemisphere (left or right). Interaction effects between factors were included. We found a significant link between post-event boundary HFO rate increases and (1) hippocampal contacts in the CA1 subfield (t=1.900, p=0.029) and (2) the interaction between right hemisphere and anterior hippocampus (t=1.912, p=0.031) (Figure 2C). This effect persists when the subject number is also included in the model as an additional covariate. We further quantified the effect of longitudinal axis and hemisphere and found a significant post-boundary hippocampal HFO rate increase in 65 right anterior hippocampal contacts (one-sample t(64)=2.828, Bonferroni-corrected p=0.006), but not in 53 left anterior hippocampal contacts (t(52)=1.291, p=0.202), 76 left posterior hippocampal contacts (t(75)=1.283, Bonferroni-corrected p=0.203), or 44 right posterior hippocampal contacts (t(43)=-0.209, Bonferroni-corrected p=0.836).

**Figure 2.**
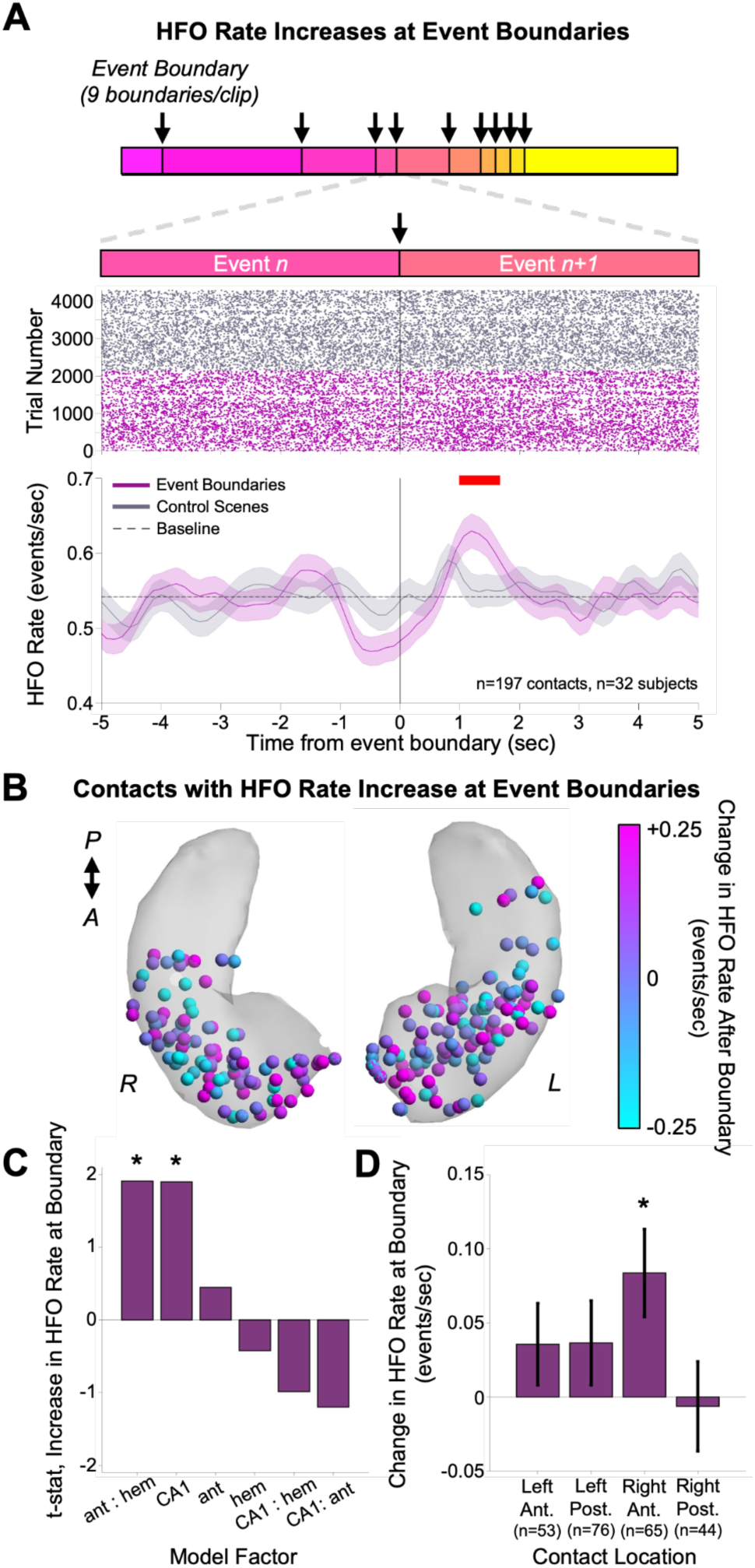
Hippocampal HFOs increase after event boundaries. (A) Hippocampal HFO rate raster plot and peri-event time histogram (PETH) time-locked to event boundaries (pink) and auditory- and visually-matched control scenes (gray) (n=197 contacts across n=32 patients, n=9 event boundaries). Red line indicates significance at p<0.05 (permutation test compared to shuffled hippocampal HFO timings in the same epoch). Shaded areas represent one bootstrap standard error of the mean computed over hippocampal HFO events.

Finally, as hippocampal HFOs may represent an electrophysiological marker for the recruitment of memory-containing regions during memory retrieval prior to onset of free recall, we created a PETH locked to the verbal recall onset of each discrete event memory. Hippocampal HFO rate increased roughly 2 seconds prior to event recall onset (n=12 patients, n=90 hippocampal contacts; p=0.043, cluster-based permutation test; Figure S5).

### Coincident hippocampal-cortical HFOs increase in regions previously linked to event segmentation

We next examined the presence of coincident HFOs during the stimulus viewing period. Since co- HFOs may coordinate the activity of hippocampal and cortical regions^21^, we hypothesized that co- HFOs are associated with event segmentation processes during the viewing of naturalistic stimuli. Our previous analysis demonstrated an increase in hippocampal HFO rate in the two-second window following event boundaries, so we examined HFOs in this window for this analysis. Using a data-driven approach, we found that co-HFOs increase after event boundaries in a wide cortical network (Bonferroni-corrected p<0.05; Figure 3A and Figures S6-S7). We hypothesized that if co-HFOs are involved in event segmentation, they should exhibit overlap with cortical regions identified in fMRI studies to subserve this process^8,29^. To quantify this, we isolated the three cortical regions identified by Hahamy et al.^29^ to be involved in event segmentation (angular gyrus, precuneus, lateral occipital cortex) and identified a significant increase in co-HFOs following event boundaries in these regions (t(109)=5.331, p<0.001, Cohen’s d=0.508; Figure S8).

**Figure 3.**
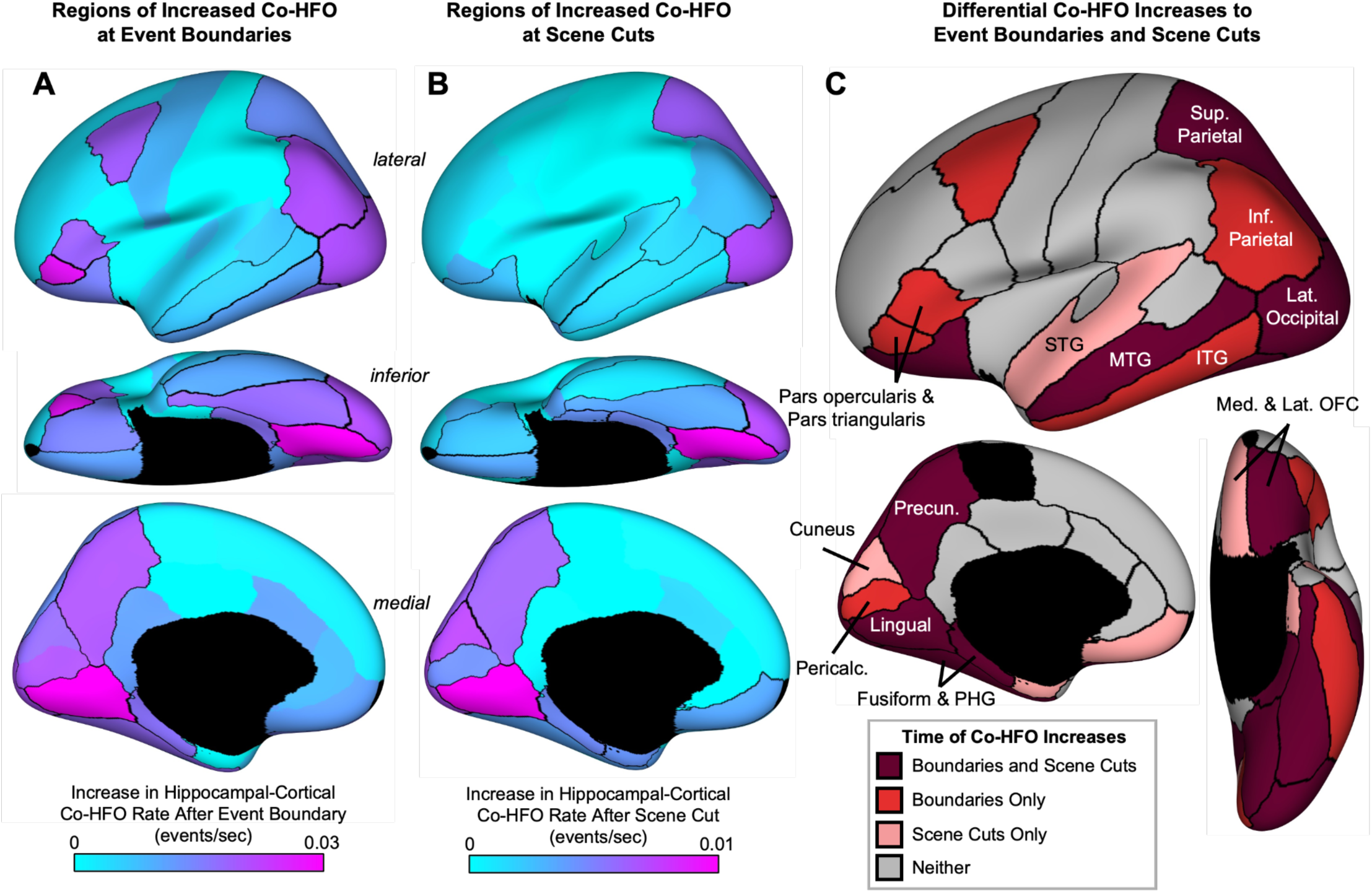
Hippocampal-cortical coincident HFOs increase in an ensemble of higher-order regions. (A) Increase in hippocampal-cortical coincident HFOs following n=9 event boundaries (compared to before boundaries) and (B) following n=145 scene cuts (compared to before cuts) averaged over all contacts within each Desikan-Killiany atlas parcel, and plotted on a normalized inflated brain (total n=32 subjects). Warmer colors indicate greater increases in coincident HFO rate. Outlined parcels denote statistical significance at p<0.05 (Bonferroni-corrected, Student’s t-test). (C) Summary plot detailing location of Desikan-Killiany atlas parcels where contacts exhibited increased coincident hippocampal-cortical HFO rate following both event boundaries and scene cuts (dark red), event boundaries only (red), scene cuts only (pink), or neither (gray). Labels are added over selected parcels to aid viewing.

We also found that co-HFOs were increased after event boundaries across both the “higher-order” (including inferior parietal cortex, lateral orbitofrontal cortex, and inferior frontal cortex) and “lower-order” (lingual gyrus, precuneus, and fusiform gyrus) event segmentation cortical regions. To determine whether co-HFOs show specificity based on the granularity of the segmentation level, we performed the same analysis as above using scene cuts (changes in camera angle or view within the continuous narrative; n=145 scene cuts within the viewed clip). Since scene cuts are known to preferentially recruit lower-order visual regions^9^, we hypothesized that co-HFOs would predominantly increase there. Indeed, we found that co-HFOs also increase after scene cuts in a wide cortical network, primarily cortical regions involved in visual processing (Bonferroni-corrected p<0.05; Figure 3B and Figure S7). Increases in co-HFO rate following event boundaries and scene cuts overlap in lower-order visual regions. Notably, the regions that exhibit increased co-HFOs after event boundaries but not scene cuts coincide with higher-order event segmentation regions that serve as association cortices (including inferior parietal cortex, inferior frontal cortex, and inferior temporal cortex) (Figure 3C).

We next aimed to link this finding with recall performance. We isolated cortical regions exhibiting increased co-HFO rate following event boundaries (Figure 3A). These regions showed higher co-HFO rate following event boundaries for events that were subsequently recalled compared to those that were not (0.035 versus 0.027 co-HFOs/sec; t(1102)=4.423, p<0.001, Cohen’s d=0.187). Notably, this relationship was not replicated for cortical regions that exhibited increased co-HFO rate following scene cuts (0.030 versus 0.029 co-HFOs/sec; t(3012)=0.619, p=0.536). Taken together, co-HFOs occur predominantly in regions that may underlie event segmentation. Further, the magnitude of co- HFOs in higher-order, but not lower-order, event segmentation regions following event boundaries relates to memory performance.

### Co-HFO motifs relate to subsequent post- boundary replay

The period immediately following an event boundary is thought to contain replay processes for the preceding event^29,31,34^. Co-HFOs may serve to synchronize brain regions involved in this process. As the content and memory representations differ across events, co-HFOs may exhibit specificity for each event. In this case, it is possible that the spatial patterns of co-HFOs are similar between event viewing and the immediately-following replay period. To investigate this, we calculated a “co-HFO index,” or a measure of the consistency in spatial and temporal co-HFO patterns between two epochs (Figure 4A). This methodology maintains spatial relationships between contacts and controls for variation in baseline co-HFO rates across contact pairs. We calculated this index by using every combination of contacts in the hippocampus and cortex. We quantified the co-HFO index between event viewing and replay (including both the immediately-succeeding replay window and all subsequent replay windows) and between event viewing and recall (Figures 4B-4D).

**Figure 4.**
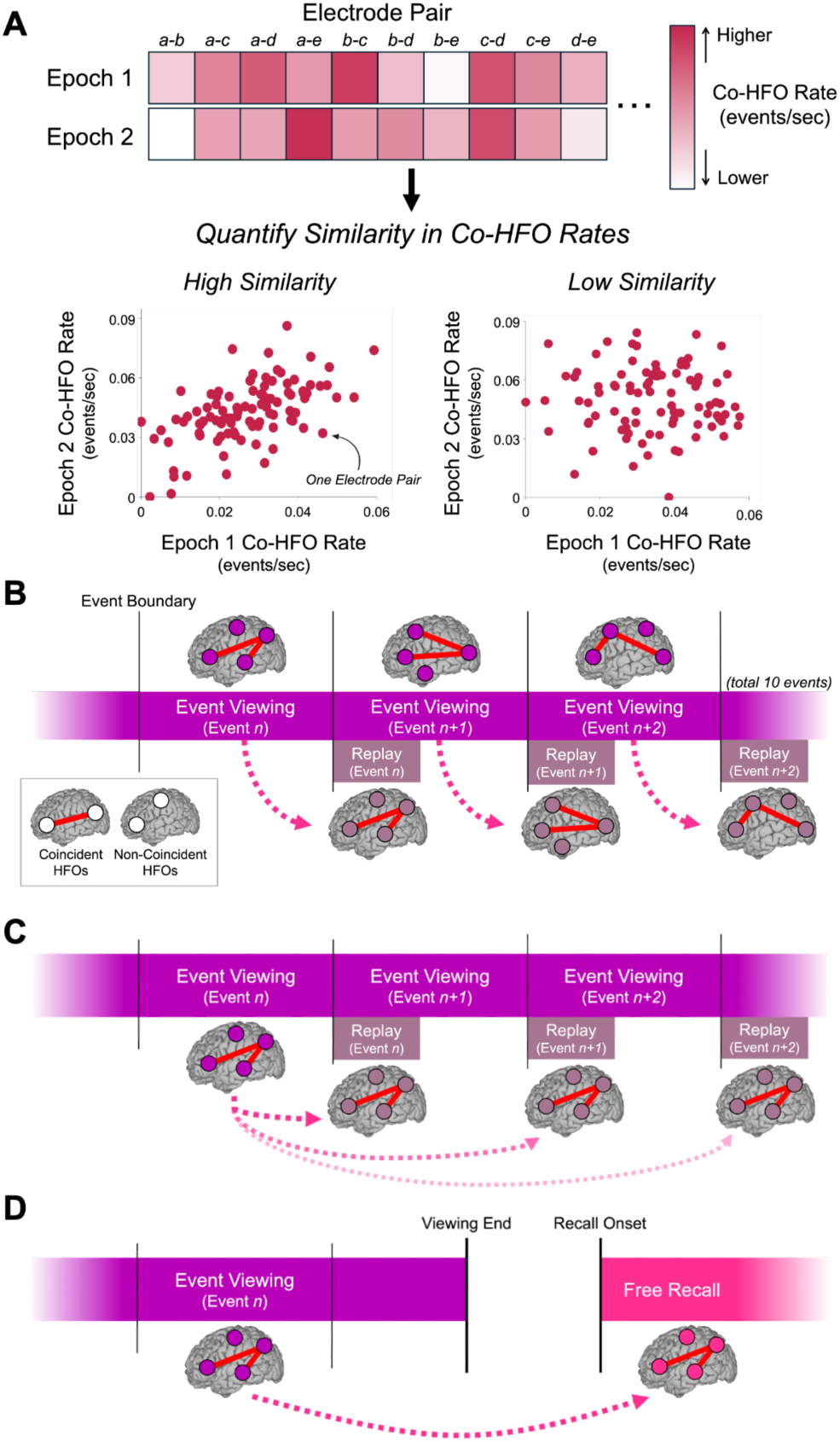
Hypotheses for co-high frequency oscillation motifs. (A) Schematic describing quantification of co-high frequency oscillation (HFO) motifs across the task paradigm. Magnitude of co-HFO pattern similarity was defined as the Pearson’s correlation coefficient of co-HFO rates for every contact pair between any two epochs (e.g. viewing and replay window). Hence, higher correlation coefficients indicate higher co-HFO motif similarity. (B-D) Tested hypotheses of co-HFO motifs: (B) co-HFO motifs may reoccur in the window immediately following event boundaries for the immediately-preceding viewed event; (C) Co-HFO motifs may reoccur in the window immediately following event boundaries for subsequent events; and (D) Co-HFO motifs may reoccur during memory retrieval.

We assessed similarity in co-HFO motifs between event viewing and replay (two-second windows following event boundaries). As we also aimed to assess subsequent replay periods, we normalized the number of events in each group by analyzing only the first five viewed events in each subject. We found high similarity in co-HFO motifs (high co-HFO index) between the event and the immediately- following replay window (p<0.001, permutation test; Hedge’s g compared to negative lag=0.377; Figure 5A). This effect is seen in 22 of 32 patients and 8 of 9 scenes (Figure S9). However, co-HFO pattern similarity continues to be significant, albeit at lower values, for the next two replay windows (p=0.023 and p=0.029; Hedge’s g=0.285 and 0.289, respectively) before diminishing (Figure 5A).

**Figure 5.**
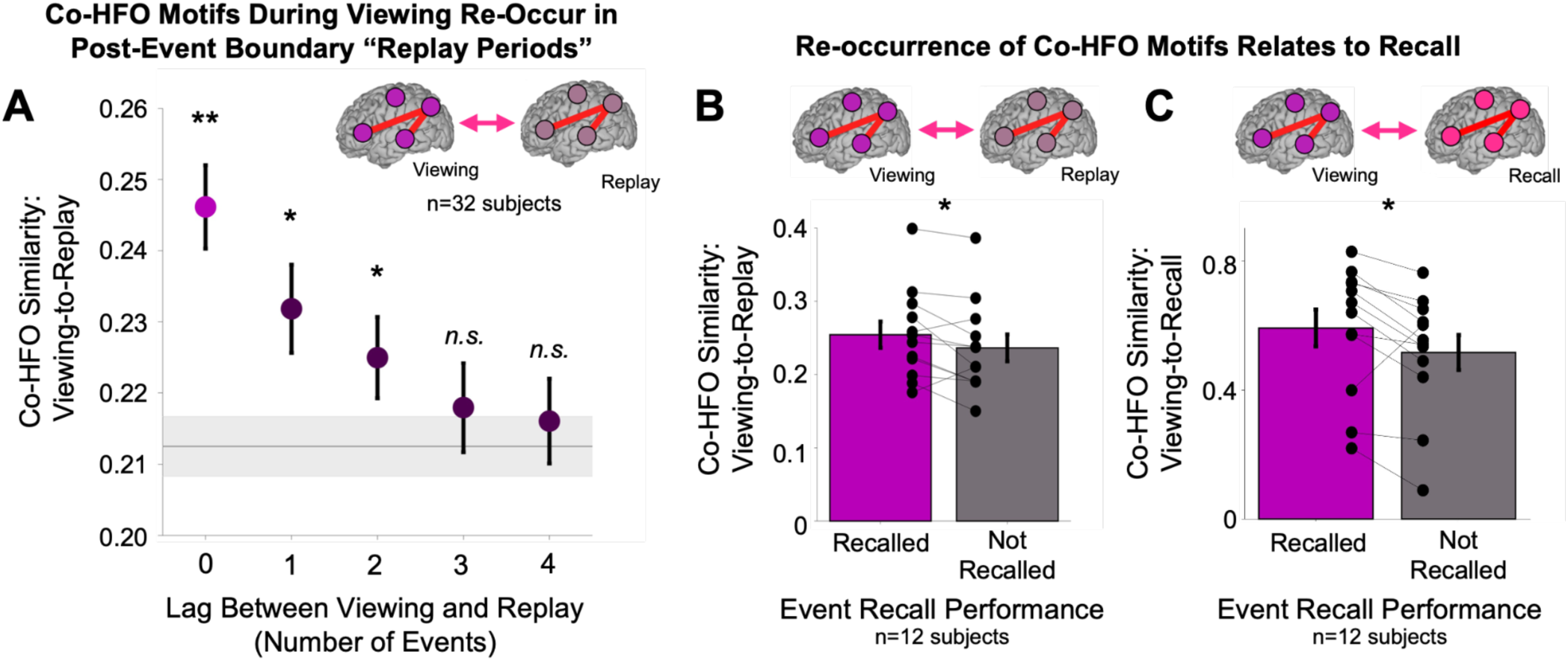
Coincident high frequency oscillation (co-HFO) motifs during viewing reoccur following event boundaries and relate to scene memory. (A) Mean similarity in co-HFO patterns for a viewed event and four subsequent post-event boundary “replay periods.” Only the first five viewed events were used in this analysis (n=32 subjects). Pink denotes the similarity between a viewed event and its immediately following post-event boundary replay period. Error bars represent one standard error of the mean. Gray horizontal line represents mean co-HFO similarity when lags are shuffled, and shaded area represents one bootstrapped standard error of the mean. ** denotes statistical significance at p<0.01 and * denotes statistical significance at p<0.05 (permutation test). (B) Group-level (n=12 subjects) mean co-HFO pattern similarity between event viewing and immediately-following replay, comparing events that were subsequently recalled or not recalled. One point represents one unique subject. * denotes statistical significance at p<0.05 (paired-samples t-test). (C) Group-level mean co-HFO pattern similarity between event viewing and free recall period, comparing events were subsequently recalled or not recalled. One point represents one unique subject. * denotes statistical significance at p<0.05 (paired-samples t-test).

Hence, co-HFO motifs show stability between scene viewing and replay, and these patterns decay as additional events are viewed and replayed. This may provide evidence for how the brain re-engages memory traces from previously-viewed scenes to encode a present one.

We then assessed the relationship between re-occurrence of co-HFO motifs and recall performance. We first assessed co-HFO similarity between scene viewing and the immediately-following replay period and found that scenes that were subsequently recalled exhibited a higher viewing-to-replay period co-HFO index compared to scenes that were not recalled (t(11)=2.248, p=0.046; Figure 5B; Hedge’s g=0.287). We then replicated this analysis between scene viewing and memory retrieval.

The co-HFO index between the recall period and the viewing of recalled events was higher compared to viewing of non-recalled events (mean 0.592±0.057 for recalled scenes compared to 0.516±0.054 for non-recalled scenes (paired t(11)=2.638, p=0.023; Hedge’s g=0.396; Figure 5C). Hence, specific patterns of co-HFOs occur during encoding that re-occur during replay of the event following event boundaries and again during memory retrieval. Furthermore, the reinstatement of these motifs relates to memory behavior.

### Hippocampal HFOs as a marker for subject-level event boundaries

Although the event boundaries that we use in this study exhibited high inter-subject consistency, it is likely that there is some variation in what each subject perceives as an event boundary. To this end, hippocampal HFOs may serve as a biomarker for “subject-specific” event boundaries. Since scenes that evoke post-viewing replay are linked to improved memory performance, and hippocampal HFOs are linked with replay processes, we investigated this by examining whether hippocampal HFO rate increases following specific scenes (as defined by scene cuts) that were later recalled by participants. We identified that there was no significant increase in hippocampal HFO rate relative to all scene cuts (Figure 6A), but when this analysis was limited to specific scenes that were later recalled, we found that hippocampal HFO rate increased 1-2 seconds following the offset of these scenes (p=0.048, cluster-based permutation test jittering HFO events; Figure 6B). Importantly, this relationship is maintained even after scenes that coincided with an event boundary were removed (8.9% of recalled scenes were within 3 seconds of an event boundary and were removed; p=0.040; Figure S10).

**Figure 6.**
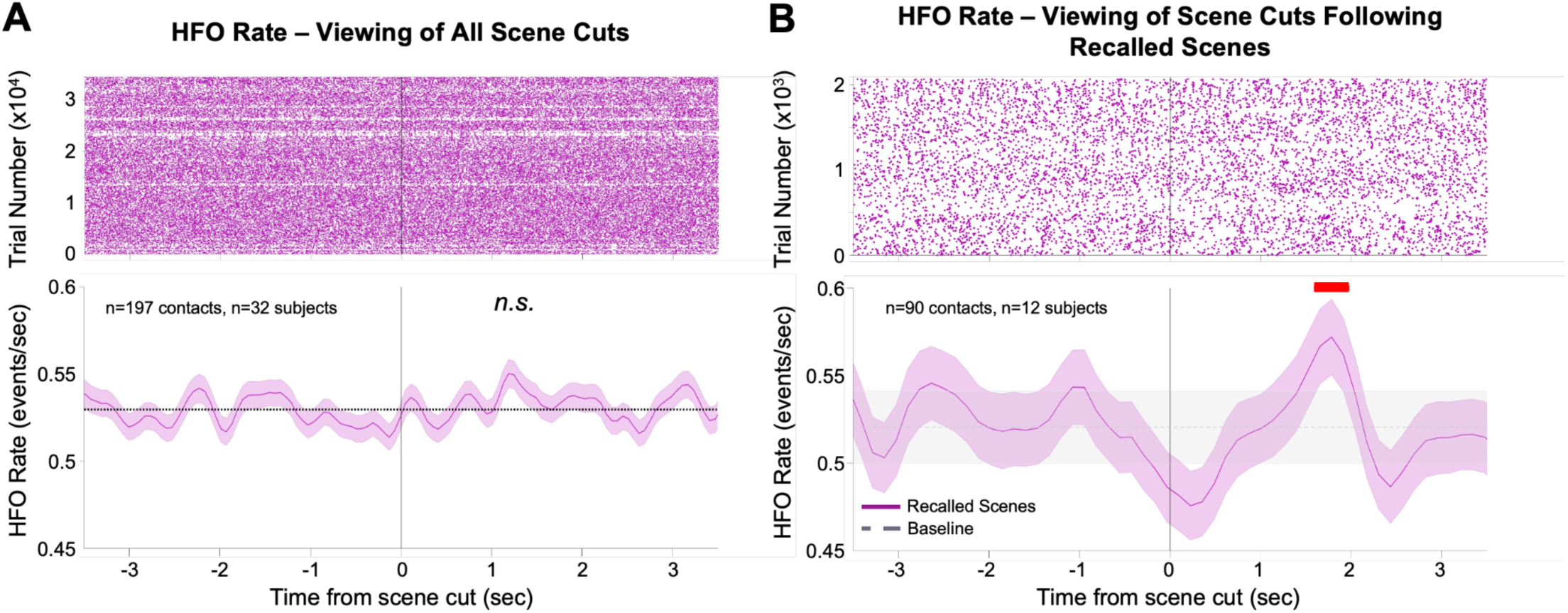
Hippocampal high frequency oscillations mark subject-specific event boundaries. (A) Hippocampal HFO rate raster plot and peri-event time histogram (PETH) time-locked to scene cuts (n=197 contacts across n=32 patients, n=144 event boundaries). Shaded areas represent one bootstrap standard error of the mean computed over hippocampal HFO events. Dotted line represents the mean hippocampal HFO rate over this epoch. n.s., not significant (permutation test compared to shuffled hippocampal HFO timings in the same epoch). (B) Hippocampal HFO rate raster plot and peri-event time histogram (PETH) time- locked to scene cuts following scenes that were recalled (n=90 contacts across n=12 patients). Shaded areas represent one bootstrap standard error of the mean computed over hippocampal HFO events. Dotted line represents the mean hippocampal HFO rate over this epoch. Red line indicates significance at p<0.05 (permutation test compared to shuffled hippocampal HFO timings in the same epoch).

## Discussion

### Summary

Experiences from the real world consist of consecutive, oftentimes interdependent, events. To process such events, two processes must occur: (1) segmentation, or chunking, of the continuous stimuli into discrete events at event boundaries; (2) consolidation of event episodic memory into cortical stores. Coincident HFOs represent a potential mechanism by which brain regions coordinate activity^21^. In this study, we examined the presence of hippocampal and cortical HFOs during the viewing and free recall of a continuous stimulus. We show that coincident HFOs in the hippocampus and specific cortical regions increase following event boundaries. The cortical regions that are recruited by co-HFOs are modulated by the granularity of the boundary. Finally, the magnitude of activation relates to subsequent memory performance. Hence, HFOs may guide event segmentation processes. Next, specific spatiotemporal patterns of co-HFOs during viewing are found to re-occur in a short window following event boundaries and during event memory retrieval. As these co-HFO motifs continue to persist several events beyond the viewed event, they may serve as a mechanism by which the brain integrates memories from previous events in the processing of subsequent ones. This is also suggestive of HFO-driven replay processes that occur following event boundaries. Taken together, human HFOs play a significant, coordinating role in the viewing, processing, and retrieval of naturalistic continuous stimuli.

### What is an HFO?

Local field potential activity can be divided into two subgroups: non-oscillatory and oscillatory. Non-oscillatory high frequency activity (also known as broadband high-frequency activity), are irregular, broad-spectrum signal fluctuations that do not follow a rhythmic pattern and are correlated with multi-unit activity^35^. High-frequency oscillatory phenomena on the other hand represent rhythmic and often regular fluctuations in neural excitability that may or may not be associated with ordered neuronal firing^36^. Coincidence of oscillations across multiple regions may lead to more optimal binding of activity, hence creating circumstances ideal for long-term potentiation^37^. Two predominant HFOs that are thought to play a significant role for memory and cognition because they relate to memory-associated neuronal firing: gamma/epsilon, and ripple^36^. Extensive studies in rodents have found that gamma/epsilon oscillations are guided by theta oscillations and predominate memory processing in the wakeful, effortful, and active states. They serve as a means of coordinating memory representations from the hippocampus to the cortex^36^. Ripple oscillations occur in offline, passive states, and serve as a biomarker for event memory replay processes^38^. However, it is perplexing that recent human studies have identified a role of gamma/epsilon oscillations during memory recall^39^ and ripple oscillations (including in a wide cortical network^21,22,25,40^) during memory encoding^18^. Further, properties of detected human ripple oscillations (rate, frequency, amplitude and duration) show stability across task states, including encoding and recall^41^. Since it is difficult to accurately disentangle different HFOs using human iEEG recordings, and HFO subtypes have very similar frequency ranges, it is possible that studies which have assessed HFOs have encompassed multiple subtypes of HFOs. We also cannot rule out that dynamics of gamma/epsilon and ripple oscillations differ in humans (for example, offline and online processes may co-occur in the awake state).

Methodological advances in HFO subtype detection may enable us to return and better elucidate the precise contribution of gamma/epsilon oscillations and ripple oscillations in our findings.

### Event segmentation and event boundaries

Event boundaries represent timepoints where a continuous narrative is segmented into discrete events, a process that is supported by a widespread cortical network^7,12^. This includes earlier hierarchical processing in regions that are more sensitive to sensory changes within the event (such as visual or auditory changes) and higher-level associative regions that are more receptive to complex or abstract changes^8,42^. fMRI and EEG studies have identified these higher-order regions to include the angular gyrus, precuneus, lateral occipital cortex, medial frontal cortex, and superior/middle temporal gyrus^8,10,14,29,42,43^. Despite this, a key question remains about how activity in these distributed nodes is coordinated, as the information stored in episodic memory represents a coherent representation integrated across time. It is theorized that this function is coordinated by the hippocampus, as hippocampal activity increases at event boundaries and is predictive of event memory^13,14,30,31,44^, and the hippocampus contains scene cells that are sensitive to event boundaries^11^. We support a function of the hippocampus in event segmentation by describing a prominent increase in HFO rate 1-2 seconds following event boundaries. This is similar to the period of hippocampal activation window that reflects replay processes^8,33^. Furthermore, hippocampal-cortical co-HFOs show specificity in location based on the granularity of the event boundary, as co- HFOs occur at higher rates in both lower-level and higher-level event segmentation regions at event boundaries, but only in lower-level visual regions at scene cuts. These findings align very closely with fMRI findings^8,29^. To extend this, we found that co-HFO rate in higher-order event segmentation regions following event boundaries relate to subsequent recall performance, a relationship that does not extend to lower-order regions. Taken together, this is supportive of a link between the hippocampus and cortical event segmentation regions that is guided by HFOs.

### Hippocampal HFOs

We leveraged this large dataset to examine specific hippocampal regions that may be more involved in coordinating activity associated with event segmentation. Specifically, we aimed to elucidate whether there were differences due to longitudinal axis, hippocampal subfield, or hemispheric laterality, as these were factors utilized by Norman et al.^17^ when highlighting left anterior hippocampal contacts within the CA1 to be most associated with autobiographical memory reactivation. Here, we show that contacts in the right anterior hippocampus and within the CA1 subfield exhibit an increase in activity at event boundaries. The hippocampal CA1 subfield is thought to play a key role in a variety of memory processes, and has been shown in rodent literature to display precise timing and spatial organization required for selective processing and replay of specific memories^38,45^. Further, hippocampal HFO events from other hippocampal subfields, such as the CA3 or dentate gyrus, do not have the same impact as the CA1 on hippocampal-cortical communication or on memory performance^46–49^. Our other findings relate to studies that indicate a specific function of the right anterior hippocampus in the longitudinal integration of memory, specifically in the utilization of prior event memories for the encoding of current, novel ones^50,51^. However, this contrasts with studies that describe a greater role of the posterior hippocampus at event boundaries^52^. To reconcile this, it is possible that hippocampal HFOs, which are most prominent in the CA1-predominant human anterior hippocampus^53^, implement differing cortical connectivity profiles when coordinating hippocampal-cortical activation^54^. Furthermore, since the timescale of hippocampal integration differs along the length of the hippocampal axis (with the anterior hippocampus integrating information along a longer timescale)^55,56^, the mechanisms underlying posterior hippocampal functional magnetic resonance imaging (MRI) activation at event boundaries may differ in function from increases in anterior hippocampal HFOs. It is also possible that HFO rate does not directly reflect increased engagement of cortical regions or networks, either because there is a basal level of HFO activity to support these functions or there are changes in HFO properties such that different HFO subtypes exhibit different functions (similar to ongoing studies investigating human theta oscillations^57,58^).

### Co-HFO index and pattern reactivation

It is thought that a rapid reactivation of event memory traces occurs at event boundaries, which functions to aid memory consolidation^59^. This process is also associated with hippocampal activation^2,8,14,15^. Further, several studies have found matching EEG^30,31^ and fMRI^29^ patterns of neural activity during event viewing and replay periods. Finally, hippocampal activation patterns appear to be unique for event memories of different types^60^. As the location and timing of co-HFOs may index specific firing patterns that encode specific memory representations^20,61^, we aimed to investigate whether spatial and temporal patterns of co-HFOs index event memory and whether such motifs re- occur following event viewing. Indeed, we found that co-HFO motifs during event viewing are unique and show similarity to the post-event replay window (two seconds following the terminating event boundary). The magnitude of co-HFO index between viewing and replay relates to subsequent event memory performance. Finally, these same motifs arise again during memory recall. This is indicative that co-HFOs across the hippocampal-cortical memory network contain event-specific representations. Co-HFO motifs may drive initial memory encoding during event viewing, and reactivate the same regions during post-event boundary replay periods and during retrieval of event memory. One interesting caveat to this approach is that it is possible that specific events engage a particular sensory modality more than others, which would affect magnitude of primary cortex co-HFO during scene viewing (e.g. a scene with vivid visual stimuli better engages the visual system during replay but not the auditory or language processing systems). To this end, it is difficult to distinguish the process of event perception, event segmentation, and memory processing; further, the extent to which primary sensory cortices contain memory representations is not well known^62^. However, it is likely that these processes are intertwined such that HFOs will reactivate the same regions that were initially recruited during stimuli presentation^18,28^.

It is thought that replay processes following event boundaries contain representations for both the immediately-preceding event and also several previous events^12,29,63–65^. This may contribute to a memory scaffold that transcends several event boundaries and binds events longitudinally. This would be key for memory of naturalistic stimuli, as despite engaging with events only once, there is an inherent narrative that may be followed and re-referenced (e.g. re-appearance of a cue from a prior event). We aimed to investigate whether co-HFO motifs persist beyond the current event and are re-referenced in future replay periods. Indeed, we found that event-specific motifs remain for three subsequent events. Importantly, as scenes became more temporally-distant, co-HFO motif similarity also decreased. This provides evidence for the reactivation of event memory from prior events for the integration of a current event into episodic memory, and that this process is facilitated by co-HFOs across the hippocampal-cortical network. Interestingly, this speaks against co-HFOs serving as a continuous scaffolding mechanism by which integrative encoding during stimulus viewing occurs^66^, as the re-occurrence of co-HFO motifs for a given event diminishes across time. However, we cannot rule out that this process instead occurs in the offline periods following the end of the entire stimulus, or that it is driven by a different mechanism.

### Who’s the driver: Hippocampus or cortex?

Co-HFOs may represent a precise means of communication between disparate areas of the brain at event boundaries. However, the directionality of this communication is not well elucidated. Initial human HFO studies examined the influence of hippocampal rippleband HFO-associated activity in task-specific cortical regions^17,18,28,67^. This aligns with prominent models of memory suggestive of hippocampal-driven influence of memory-containing cortical regions during encoding and recall^38^.

However, more recent studies have identified a cortico-cortical pattern of influence^21,22,25^. This may more closely align with theories proposing a significant role of cortical regions in influencing other cortical regions and the hippocampus^68^. In our study, we found both hippocampal-cortical and cortico-cortical co-HFO dynamics around timepoints thought to be relevant for episodic memory function.

Although we were not able to precisely determine directionality due to the limited spatial and temporal sampling of our technique, future studies should consider the possibility that select cortical regions may be driving synchronized memory network activity (which may be in conjunction with, or independent of, the hippocampus).

### Subject-specific event boundaries

We also extended our knowledge of event segmentation to “subject-specific event boundaries.” Thus far, studies have utilized event boundaries that have a high inter-subject consistency^69^, including this study^9^. Although there is some objectivity in this approach of assigning event boundaries, it inherently leads to potential inter-subject variation. Though this variation may be mitigated by large datasets, there is also value in assessing such differences. Here, we assessed whether hippocampal HFOs may serve as a biomarker for subject-level event boundaries. The rationale for this was based on prior studies that identified scene chunks associated with event segmentation to be better encoded and have a higher likelihood of being subsequently recalled^70,71^. To investigate this, we divided the stimuli into more granular segments using scene cuts (as they have clear onset and offsets) and found increases in hippocampal HFO rate 1-2 seconds following scene cut offset for scenes that were later recalled but not for scenes that were not later recalled. This relationship is maintained even after the removal of scene cuts that coincide with event boundaries. Hence, it is possible that hippocampal HFOs are an electrophysiological marker for subject-specific event boundaries. Future studies should investigate discrepancies between population-level and subject-specific event boundaries.

## Conclusions

Coincident hippocampal and cortical HFOs may enable the brain to process and store naturalistic stimuli by coordinating the networks underlying event segmentation and episodic memory processing. Event-specific spatiotemporal patterns of coincident HFOs exist during scene viewing that are replayed immediately following event boundaries and again during retrieval of scene memory. Hence, HFOs represent a robust mechanism that drives the encoding and retrieval of rich, continuous episodic memories from our everyday lives.

## Methods

### Subjects

Intracranial recordings were obtained from 30 patients (14 females; 32 testing sessions) with medically-intractable epilepsy undergoing iEEG recording at Northwell Health (New York, USA) to identify epileptogenic zones for potential surgical treatment (Table S1). Data from a subset of these participants (n=23) was used in a previous study^9^. These subjects are continuously monitored with intracranial contacts that were stereotactically-placed, subdural grid or strip contacts, or both, for a period of up to 4 weeks. During this time, they may participate in cognitive and functional testing. The decision to implant, location of implanted contacts, and duration of implantation were made exclusively on clinical grounds by the clinical treatment team. Subjects were invited to participate in this study if their iEEG recording included a hippocampal contact and ability to maintain attention to the 10-minute audiovisual stimuli task. This study was conducted in accordance with the Institutional Review Board at the Feinstein Institutes for Medical Research (Northwell Health), and informed consent was obtained prior to research testing. No clinical seizures occurred during or within the two-hour period prior to the experimental block. All participants performed the task in English.

### Stimuli and Task

Psychophysics Toolbox (version 2014-10-19_V3; Gstreamer version 1.10.2)^72^ running on MATLAB R2012b (Mathworks, Natick, MA) was utilized for precise stimuli presentation. Patients viewed 600 seconds of the animated feature film “Despicable Me” without any explicit memory instruction. The audiovisual clip was presented continuously. Nentwich et al. previously identified 9 event boundaries and 145 scene cuts (a change in viewing angle or scene; mean inter-cut interval 4.6 seconds) in this clip, which we utilized for analysis. We also implemented 9 scenes that were matched (audio and visual) to event boundaries to control for any inherent properties of event boundaries (e.g. changes in attention or movement) that may impact analysis^9^.

Immediately following viewing, participants were verbally asked to “describe, in as many details as you can, as many things as you can recall from the movie.”. Patients verbally indicated when they were finished with their recall. The verbal responses were recorded by a microphone affixed to a nearby stable surface for further assessment and quantification in offline analysis. Offline, the onsets, offsets, and content of each recall were extracted in an offline analysis using Audacity auditory presentation software (Audacity, Oak Park, MI, USA). This included the onset of recall for each unique event memory representation or specific, differing aspect of an event. For each viewed scene (as segmented using scene cuts), it was identified whether the subject recalled the scene during recall.

Notably, since subjects varied in the number of contacts and recall performance, all analysis relating electrophysiological activity during encoding to memory performance was performed on the group level (n=12 subjects).

### Intracranial Recordings

Intracranial recording sites were stereoelectroencephalography depth electrodes, subdural grids, and/or subdural strips (Ad-Tech Medical Instrument Corp., Oak Creek, WI, USA; Integra LifeSciences, Princeton, NJ, USA; PMT Corp., Chanhassen, MN). Subdural grids/strip contacts were 3-mm platinum disks with 10-mm intercontact spacing. Depth electrode contacts were 2mm cylinders with 0.8mm diameter and 4.4mm or 2.2mm intercontact spacing. During the recordings, the intracranial electrode signal was referenced to a subdermal electrode or subdural strip. Neural signals were acquired using a Tucker-Davis Technologies (TDT) PZ5M module (Tucker-Davis Technologies Inc., Alachua, FL) at either 500Hz, 1.5kHz, or 3kHz. During the task, transistor-transistor logic pulses were utilized to align onset and offset of audiovisual clip to synchronize stimuli presentation with neural data.

Data analysis was performed in MATLAB R2023b using Fieldtrip^73^ and custom analysis scripts. Data was resampled to 500Hz. Power-line noise was removed using a notch filter (zero-lag linear-phase Hanning-window FIR bandstop filter) at 60Hz, 120Hz, and 180Hz. Raw iEEG data was inspected visually to detect noisy or bad channels, which were removed from all subsequent analysis. Channels that were identified as being within the seizure onset zone were excluded from analysis. Data was average-referenced to remove global artifact.

As study participants underwent clinical care alongside research testing, we implemented notes derived from clinical iEEG review (performed or supervised by author SB) to exclude all contacts that exhibited any electrophysiological signs of epileptic pathology. This included contacts that exhibited IEDs or were closely related to epileptic foci.

### Hippocampal and Cortical Contact Localization

Prior to electrode implantation, a T1w magnetic resonance imaging (MRI) scan was performed. After implantation, a CT scan was performed. We utilized the iElvis toolbox^74^, Bioimage suite^75^, and Freesurfer^76^ for intracranial electrode localization. Briefly, electrodes were manually registered to the post-implantation CT scan using BioImage Suite, which was co-registered to the pre-implantation MRI scan. These images were converted to a standard coordinate space, the cortical and hippocampal fields were segmented, and anatomical locations for each segment was assigned. Finally, contact locations were projected onto the standardized brain space. For all anatomical region of interest analysis, these parcellations were utilized to select contacts of interest.

### Hippocampal and Cortical HFO Detection

HFO detection was performed similar to prior studies^17,18,28^ and with close consideration of established protocols for HFO detection^53^. The following procedure was repeated for each contact.

For each hippocampal and cortical contact, the signal was bipolar-referenced to a nearby white matter contact, defined as a contact containing a proximal tissue density value of less than -0.9^77^. This aimed to optimize signal by reducing noise. The resulting re-referenced signal was filtered between 80 and 140Hz using a zero-lag, linear-phase Hanning window finite impulse response filter (5Hz transition band). A Hilbert transform was utilized to attain HFOband power envelope. The signal was clipped to 3 standard deviations, then squared and smoothed using a Kaiser-window finite impulse response low-pass filter with 40Hz cutoff. The mean and standard deviation of the entire audiovisual clip period was implemented to attain a baseline for event detection. Events from the power envelope that exceeded 3 standard deviations above baseline were selected as potential HFO events. The onset and offset of each event was defined as the timepoints where the 80-140Hz power decreased below 2 standard deviations above baseline. Events shorter than 42ms (computed as duration of 3 cycles of 70Hz) and longer than 250ms were removed. Events where the peak-to-peak duration was under 200ms were merged. HFO peak was then aligned to the nearest maxima of the bipolar-referenced trace.

To control for artifacts, a control detection was executed on the common average signal of all contacts. Any HFO events that occurred within 50ms of a common-average 80-140Hz peak were removed. As pathologic discharge events (interictal epileptic discharges; IEDs) may appear similar to HFOs, we implemented a stringent automated detection process for their removal. Each bipolar-referenced trace was filtered between 25 and 60Hz (using a zero-lag linear-phase Hamming window finite impulse recovery filter) and a similar methodology to above was implemented (Hilbert transform, square, normalize). Detected events that exceeded 5 standard deviations were marked as IEDs, and all HFO events occurring within 200ms of these events were excluded. To confirm that all detected HFOs were oscillatory, we implemented the eBOSC toolbox^78^ and confirmed that each HFO event had at least one oscillatory cycle within the 80-140Hz frequency range.

Properties of each HFO were then extracted, including: HFO duration (onset-to-offset), HFO amplitude (from the bipolar-referenced trace), and peak frequency (as provided by the eBOSC toolbox).

### Peri-event time histograms

To construct peri-event time histograms of HFO rates locked to specific timepoints of interest, we identified and used a bin width advised by Scott’s optimization method, which optimizes histogram bin size to event density^79^ (Scott; 1977). We typically implemented 3- or 4-point smoothing to aid in visualization. To determine significant timebins, 2000 iterations of peri-event time histograms were computed by circularly jittering HFO times across the entire peri-event epoch of interest. Cluster-based permutation tests were implemented to determine timebins with significant increase in HFO rate.

### Coincident HFO events

A coincident HFO event was defined as two HFO events where the peak-to-peak duration was less than 100ms (defined as half of the maximum 200ms duration). This was selected because HFO peaks are more readily detectable than the onset and offset. Rates of coincidence were calculated between the hippocampus and all cortical sites, and between each pair of cortical sites. Changes in the rate, for example around event boundaries or scene cuts, were calculated in coincident events per second.

### Quantification of Co-HFO Motifs

We next aimed to quantify similarity in co-HFOs across the entire brain (all contacts) during encoding, replay, and retrieval. Hence, we developed a methodology that enabled quantification of co-HFO similarity between any two time periods using the rate of co-HFOs in each epoch. Notably, this methodology maintains spatial relationships between contacts, is specific for each event (or time window), and mitigates any baseline co-HFO differences between contact pairs.

We quantify co-HFO index for each subject. For each contact pair within this subject, we quantify the rate of coincident HFOs for each time window of interest. For each epoch of interest, we get a vector where each contact pair contributes one value. Then, for any given two time epochs, we perform a correlation between these vectors. The resulting correlation coefficient is the co-HFO index. In this way, a high co-HFO index indicates that the magnitude of coincident HFOs was more similar between the two epochs of interest, and a low co-HFO index indicates that the magnitude of coincident HFOs was less similar between the two time windows. Critically, this approach does not necessarily evaluate the contribution of coincident HFOs between any two given contact pairs, but rather uses a “whole brain” perspective where HFOs are relatively ubiquitous across the cortex and fluctuate in rates and coincidence. The significance of the derived co-HFO index values were calculated using a permutation test where the HFO timings of coincident HFOs were jittered in the analysis window.

To ensure that the witnessed effects were not driven by any specific contact pair or pairs of regions, we then computed co-HFO indices between scene viewing and the immediately-following replay window where one pair of contacts were removed from analysis. The change in co-HFO index value was taken to reflect the magnitude of effect that this pair of contacts had on this analysis. We then repeated this process for all contact pairs. To visualize this data, we created a matrix where the magnitude of effect for contacts in each pair of cortical parcels was plotted. We did not identify any regions that visually appear to make a significant impact on the co-HFO index (Figure S11).

### General linear model

A general linear model was utilized to investigate specific properties of contacts that contributed most to the increase in HFO rate following event boundaries. The variables selected for analysis (contact in CA1 versus not, contact in anterior versus posterior hippocampus, and left versus right hemisphere) were selected based on a prior study that utilized the same factors (Norman 2021). We assessed the impact of each factor and all interaction effects on the magnitude of increase in HFO rate following event boundaries. This normalized our analysis to account for any variation in baseline HFO rate across contacts. To assess for significance, we generated a permutation distribution for the beta-value of each factor and interaction effect by shuffling contact labels 2000 times and recalculating the model for each iteration. To confirm that the effects depicted in our analysis are also seen on a non-modeled dataset, we also depict the increase in HFO rate following event boundaries by hemisphere and longitudinal position (Figure 2) and by hippocampal subfield (Figure S13).

HFO rate increase ∼ CA1 + LongitudinalPosition + Hemisphere + CA1*LongitudinalPosition + CA1*Hemisphere + LongitudinalPosition*Hemisphere + CA1*Longitudinal Position*Hemisphere

## Acknowledgements

We would like to thank the patients for volunteering their time and effort to participate in our study. We also thank the Epilepsy Monitoring Staff at the North Shore University Hospital and Lenox Hill Hospital for their support.

## Authorship Statement

AM, GT, MN, and SB designed the study. ADM performed clinical electrode implantation. AM, GT, MN, EE, NM, JW, EF, and SG conducted the experiments. AM, GT, and NM conducted all data analysis. AM and SB prepared the initial draft of the manuscript. All authors approved of the final manuscript draft.

