## SupplementaryMaterial for "Motifs of human hippocampal and cortical high frequency oscillations structure processing and memory of naturalistic stimuli"

### Supplementary Information

**Table 1.** Demographics of participants.

| Patient Number | Age | Sex | Recall Task? | # of Hippocampal Contacts |
|---|---|---|---|---|
| 1 | 49 | M | No | 2 |
| 2 | 36 | F | No | 6 |
| 3 | 46 | M | No | 16 |
| 4 | 29 | F | No | 13 |
| 5 | 28 | F | No | 8 |
| 6 | 44 | F | No | 8 |
| 7 | 37 | M | No | 3 |
| 8 | 20 | M | No | 9 |
| 9 | 55 | M | No | 4 |
| 10 | 53 | F | No | 7 |
| 11 | 53 | M | No | 8 |
| 12 | 52 | M | No | 11 |
| 13 | 23 | M | No | 6 |
| 14 | 40 | M | No | 9 |
| 15 | 25 | M | No | 12 |
| 16 | 51 | F | No | 3 |
| 17 | 43 | M | No | 4 |
| 18 | 36 | F | No | 9 |
| 19 | 48 | M | No | 4 |
| 20 | 24 | M | No | 6 |
| 21 | 33 | M | Yes | 5 |
| 22 | 61 | M | Yes | 5 |
| 23 | 22 | M | Yes | 9 |
| 24 | 46 | F | Yes | 7 |
| 25 | 60 | F | Yes | 7 |
| 26 | 37 | F | Yes | 4 |
| 27 | 37 | F | Yes | 13 |
| 28 | 20 | M | Yes | 5 |
| 29 | 42 | M | Yes | 6 |
| 30 | 33 | F | Yes | 10 |
| 31 | 56 | F | Yes | 8 |
| 32 | 27 | F | Yes | 11 |

**Table 2.** Number of contacts for each DK parcel across all subjects, and number of subjects with contacts implanted in each DK parcel.

| DK Parcel Name | #contacts | #subjects |
|---|---|---|
| bankssts | 14 | 7 |
| caudalanteriorcingulate | 30 | 11 |
| caudalmiddlefrontal | 56 | 18 |
| cuneus | 30 | 8 |
| entorhinal | 43 | 18 |
| frontalpole | 6 | 2 |
| fusiform | 162 | 29 |
| inferiorparietal | 49 | 19 |
| inferiortemporal | 193 | 28 |
| insula | 260 | 29 |
| isthmuscingulate | 28 | 15 |
| lateraloccipital | 58 | 11 |
| lateralorbitofrontal | 203 | 30 |
| lingual | 33 | 8 |
| medialorbitofrontal | 102 | 27 |
| middletemporal | 284 | 31 |
| paracentral | 25 | 13 |
| parahippocampal | 74 | 27 |
| parsopercularis | 96 | 27 |
| parsorbitalis | 41 | 15 |
| parstriangularis | 87 | 29 |
| pericalcarine | 7 | 2 |
| postcentral | 75 | 22 |
| posteriorcingulate | 42 | 13 |
| precentral | 97 | 27 |
| precuneus | 70 | 20 |
| rostralanteriorcingulate | 62 | 20 |
| rostralmiddlefrontal | 125 | 23 |
| superiorfrontal | 151 | 24 |
| superiorparietal | 60 | 14 |
| superiortemporal | 364 | 32 |
| supramarginal | 104 | 25 |
| temporalpole | 32 | 15 |
| transversetemporal | 32 | 15 |

**Figure S1.** Number of contacts across n=32 subjects for each cortical parcel (Desikan-Killiany atlas).

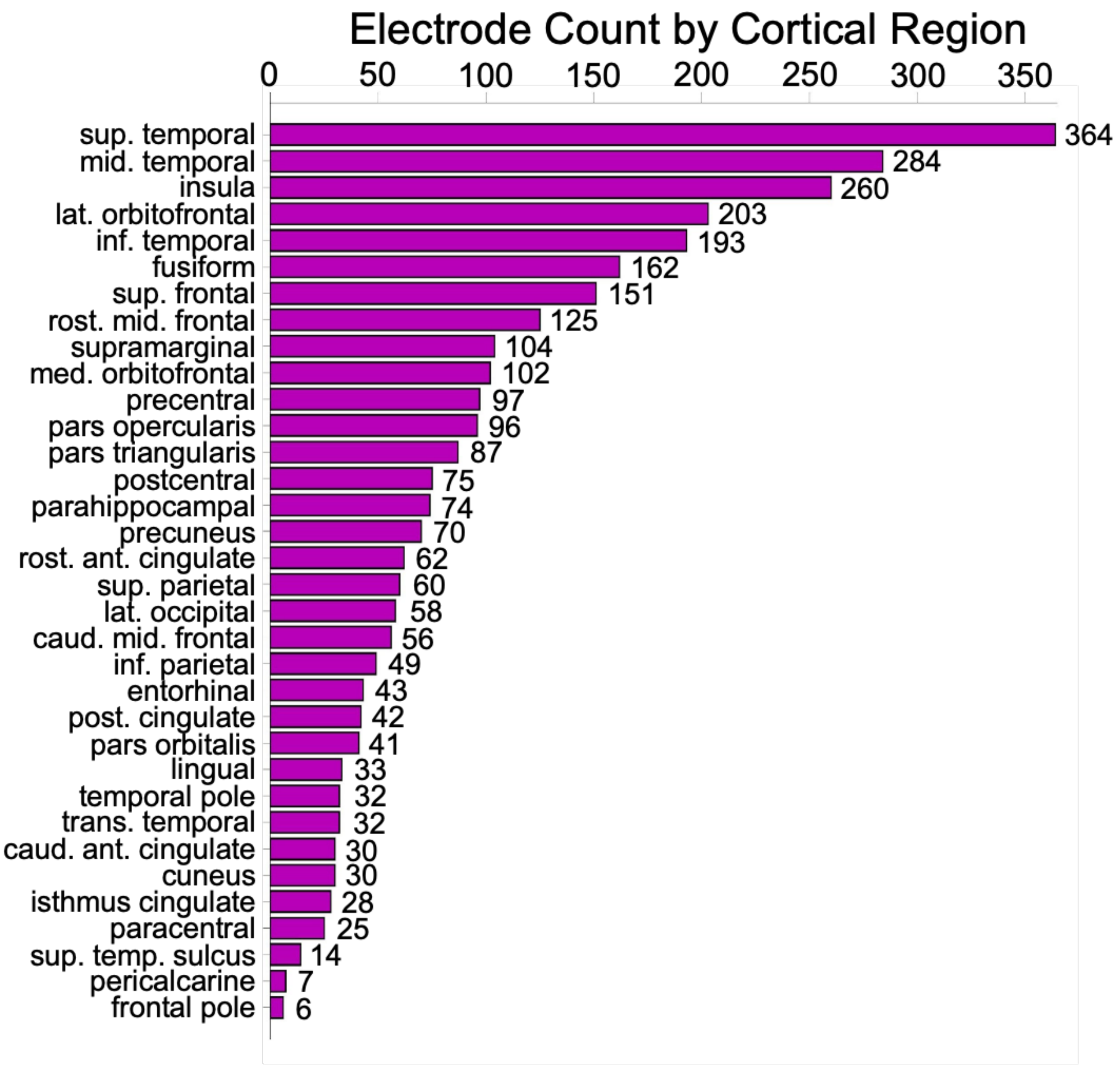

**Figure S2**. (Top) Overall rate, (middle) mean peak frequency, and (bottom) mean duration for hippocampal and cortical HFOs during movie viewing (n=32 subjects). Non-hippocampal contacts are divided into cortical parcels using the Desikan-Killiany atlas. Error bars represent one standard error of the mean, computed across contacts.

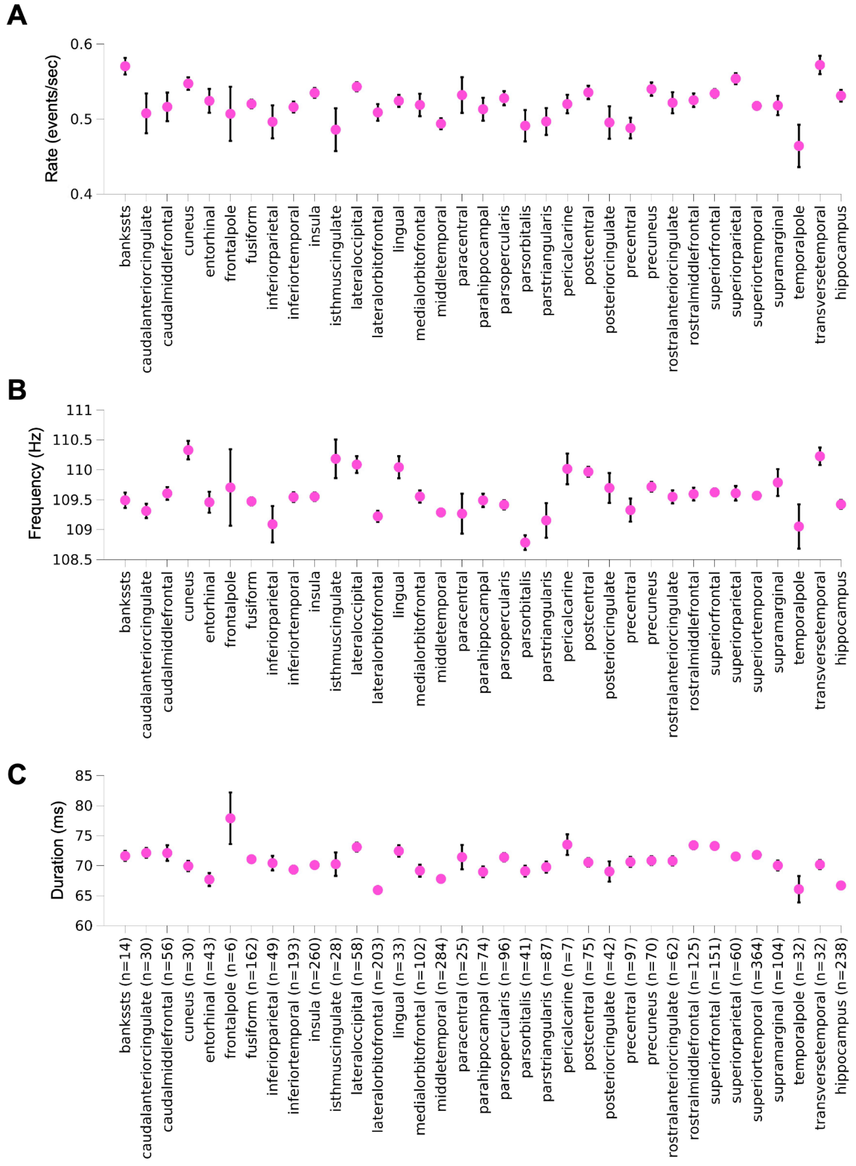

**Figure S3**. Amplitude of noise envelope across n=9 event boundaries (red) and n=9 auditory-matched control scenes (black). Shaded areas represent standard error of the mean computed across boundaries or control scenes. Thin dotted lines represent the raw auditory envelope used to compute the mean. N.s., not significant.

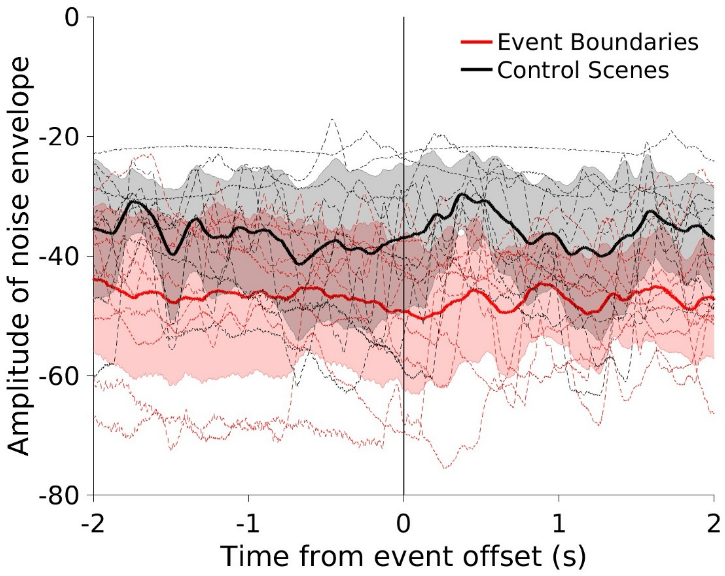

**Figure S4.** Bar graph comparing hippocampal HFO rate before and after event boundaries on the group level. Mean represents hippocampal HFO rate in the two-second window before event boundaries (light purple) and in the two-second window after event boundaries (dark purple) across subjects. Error bars represent standard error of the mean. Each pair of points that are connected by a line represent values from each individual subject.

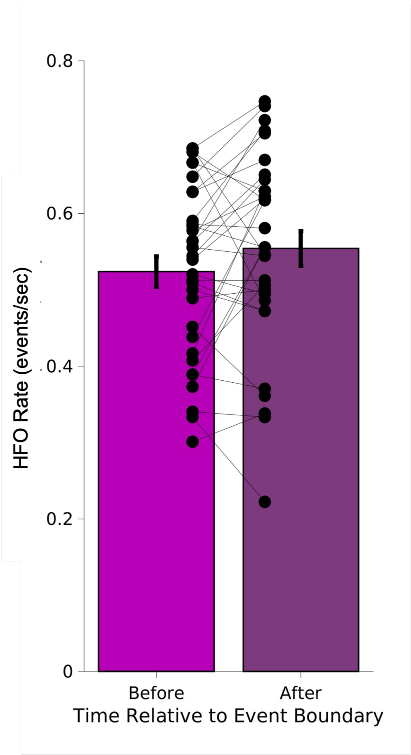

**Figure S5.** Hippocampal HFO rate raster plot (top) and peri-event time histogram (below) time-locked to onset of verbal recall of a unique scene. Red line indicates significance at $p<0.05$ (permutation test compared to shuffled hippocampal HFO timings in the same epoch). Shaded areas represent one bootstrap standard error of the mean computed over hippocampal HFO events. Dotted line represents the mean hippocampal HFO rate over this epoch.

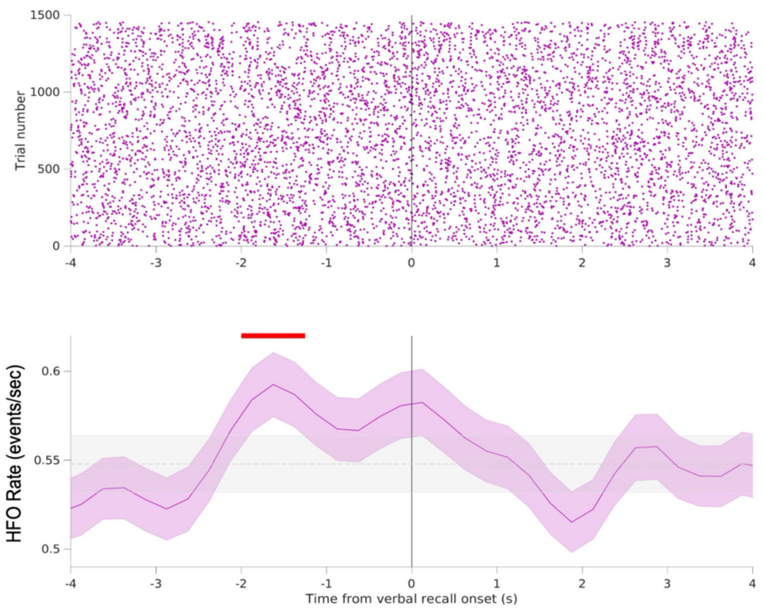

**Figure S6.** Rate of coincident HFOs during viewing between each cortical parcel, averaged across patients, contact pairs, and hemisphere. Contacts were divided into cortical regions using the Desikan-Killiany atlas. Cortical regions are sorted by brain lobe. Warmer values reflect increased magnitude of coincident HFOs.

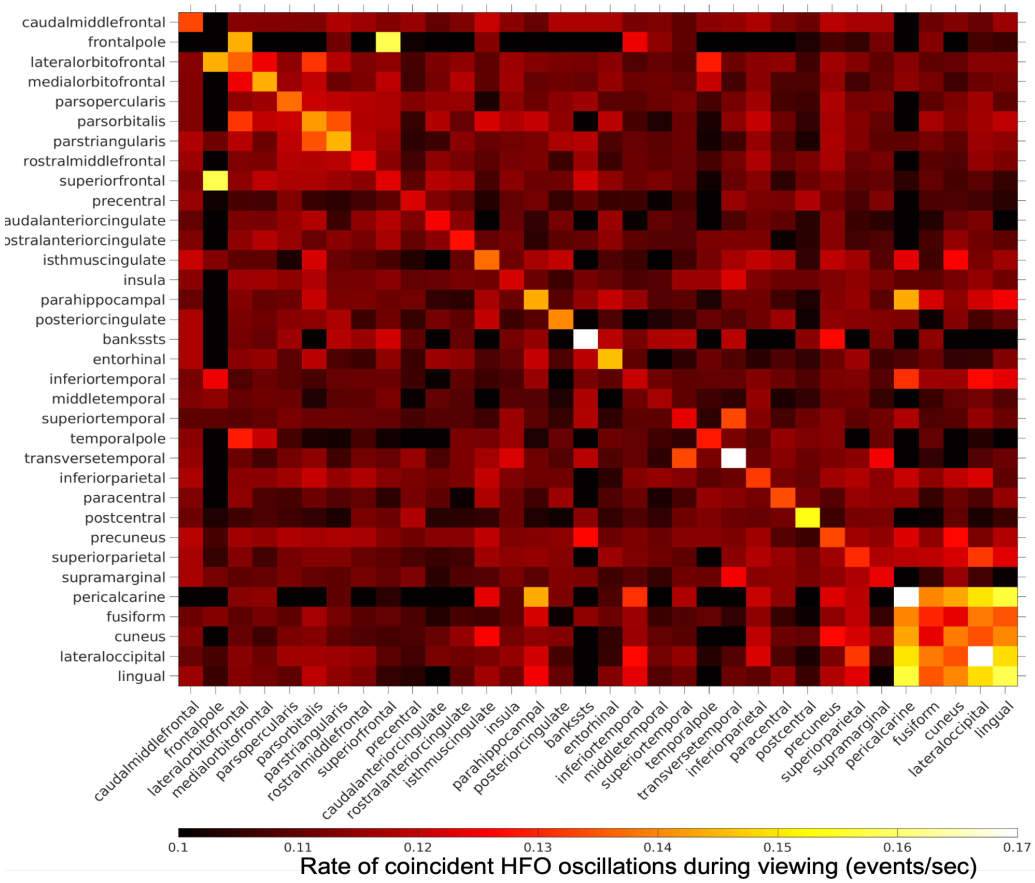

**Figure S7.** Magnitude of increase in hippocampal-cortical co-HFO rate following (A) event boundaries and (B) scene cuts across n=32 subjects. Contacts were divided into cortical regions using the Desikan-Killiany atlas. Error bars represent one standard error of the mean computed across cortical-hippocampal contact pairs. Red stars denote significance at p<0.05 (Bonferroni-corrected).

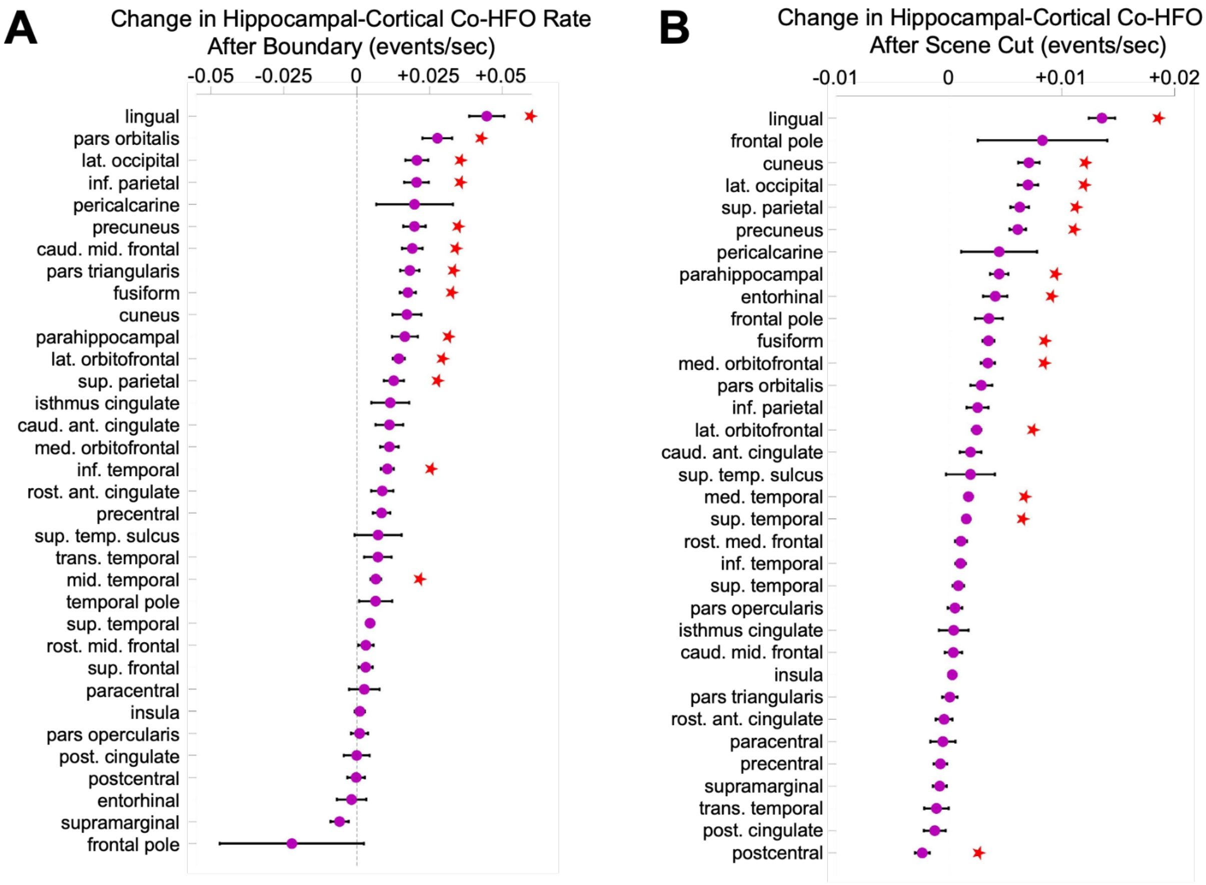

**Figure S8.** Increase in co-HFO rate following event boundaries for the three cortical regions thought to play a role at event boundaries as identified by Hahamy et al.[29]. Error bars represent one standard error of the mean computed across all cortical-hippocampal contact pairs. *** represents statistical significance at $p<0.001$.

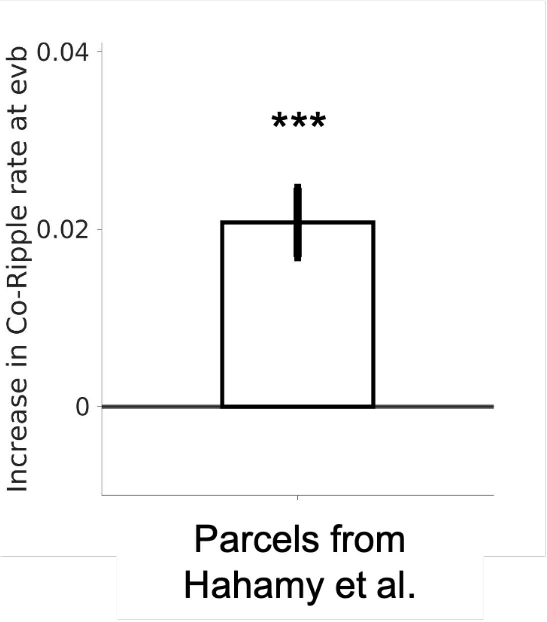

**Figure S9.** Mean co-HFO index calculated between event viewing to the two-second "replay window" following event boundaries, calculated for (A) each subject and (B) each event. Distributions represent results of a permutation test for each subject or event calculated by randomly jittering HFO timing for each contact, calculating a new co-HFO index value, and repeating this procedure 2000 times. Statistical significance at the $p<0.05$ level was computed by comparing the real value with the permutation distributions, and was found for 22 of 32 subjects (excluding subjects 2, 5, 8, 10, 16, 24, 25, 26, 27, 32) and 8 of 9 events (excluding event 2).

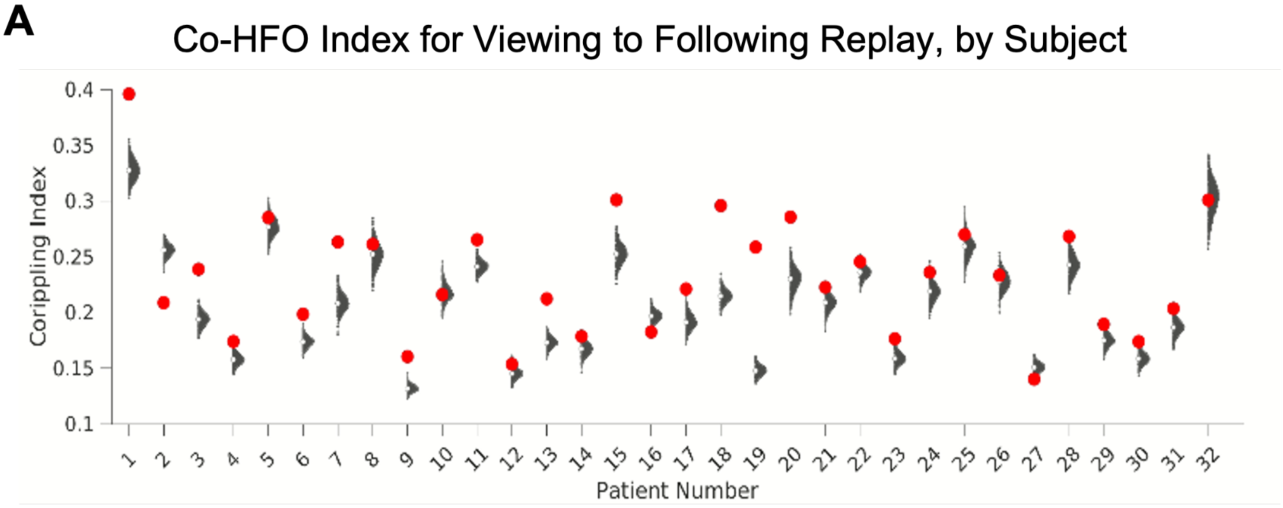


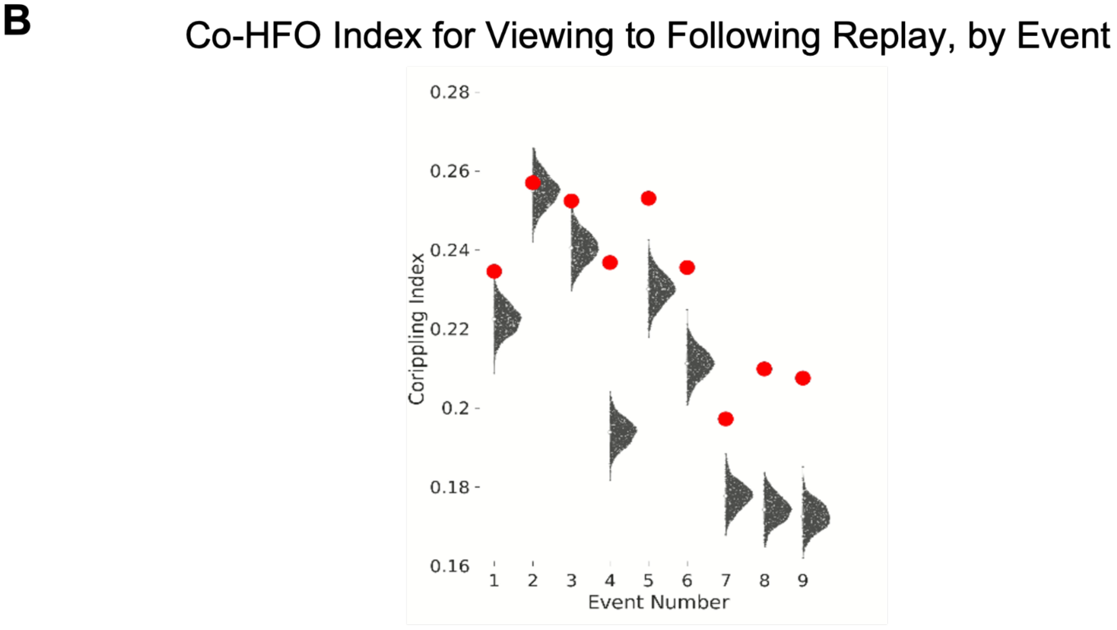

**Figure S10.** Hippocampal HFO rate raster plot (top) and peri-event time histogram (below) time-locked to the offset of scenes that were subsequently recalled. Scenes that were within 3 seconds of an event boundary were excluded from this analysis. Red line indicates significance at $p<0.05$ (permutation test compared to shuffled hippocampal HFO timings in the same epoch). Shaded areas represent one bootstrap standard error of the mean computed over hippocampal HFO events. Dotted line represents the mean hippocampal HFO rate over this epoch.

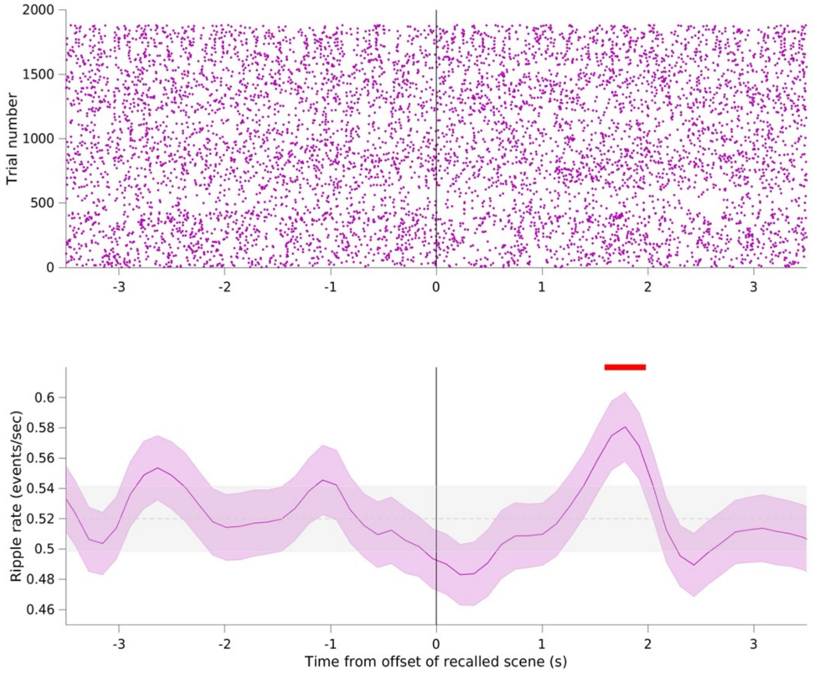

**Figure S11.** Contact pairs contributing to event viewing-to-replay co-HFO index. Cortical contacts are divided by cortical region using the Desikan-Killiany atlas. Values are calculated by removing each contact pair, re-calculating co-HFO index, and performing this analysis for all contact pairs across all patients. Negative values indicate that the pair more significantly contributed to co-HFO index values. Values are normalized for ease of visualization.

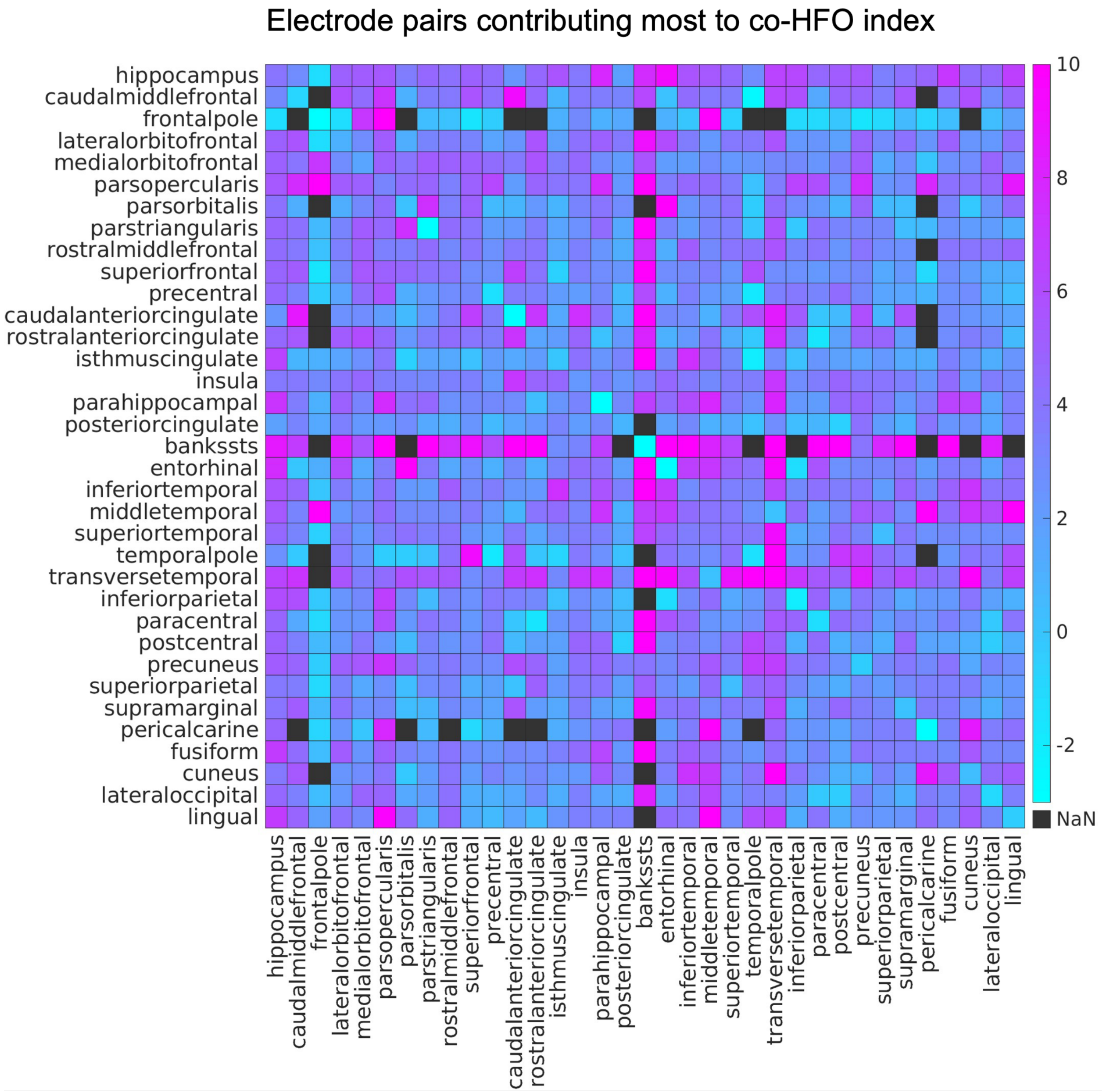
